# High resolution mapping of a *Hordeum bulbosum*-derived powdery mildew resistance locus in barley using distinct homologous introgression lines

**DOI:** 10.1101/867960

**Authors:** Parastoo Hoseinzadeh, Brigitte Ruge-Wehling, Patrick Schweizer, Nils Stein, Hélène Pidon

## Abstract

Powdery mildew caused by *Blumeria graminis* f.sp. *hordei (Bgh)* is one of the main foliar diseases in barley (*Hordeum vulgare* L.; *Hv*). Naturally occurring resistance genes used in barley breeding are a cost effective and environmentally sustainable strategy to minimize the impact of pathogens, however, the primary gene pool of *H. vulgare* contains limited diversity owing to recent domestication bottlenecks. To ensure durable resistance against this pathogen, more genes are required that could be unraveled by investigation of secondary barley gene-pool. A large set of *Hordeum bulbosum* (*Hb*) introgression lines (ILs) harboring a diverse set of desirable resistance traits have been developed and are being routinely used as source of novel diversity in gene mapping studies. Nevertheless, this strategy is often compromised by a lack of recombination between the introgressed fragment and the orthologous chromosome of the barley genome. In this study, we fine-mapped a *Hb* gene conferring resistance to barley powdery mildew. The initial genotyping of two *Hb* ILs mapping populations with differently sized 2HS introgressions revealed severely reduced interspecific recombination in the region of the introgressed segment, preventing precise localization of the gene. To overcome this difficulty, we developed an alternative strategy, exploiting intraspecific recombination by crossing two *Hv/Hb* ILs with collinear *Hb* introgressions, one of which carries a powdery mildew resistance gene, while the other doesn’t. The intraspecific recombination rate in the *Hb*-introgressed fragment of 2HS was approximately 20 times higher than it was in the initial simple ILs mapping populations. Using high-throughput genotyping-by-sequencing (GBS), we allocated the resistance gene to a 1.4 Mb interval, based on an estimate using the *Hv* genome as reference, in populations of only 103 and 146 individuals respectively, similar to what is expected at this locus in barley. The most likely candidate resistance gene within this interval encodes a legume-type lectin-receptor-like protein (LecRLP). Like other LecRLPs that have been implicated in resistance, this gene could be a good candidate for *Hb* resistance. The reported strategy can be applied as a general strategic approach for identifying genes underlying traits of interest in crop wild relatives.

## 1 Introduction

The increased genetic uniformity of cultivated crops, makes them highly vulnerable to various biotic and abiotic stresses, leading to crop yield losses and serious food security issues (Hoisington et al., 1999). Disease resistance breeding is a cost effective and environmentally sustainable strategy for minimizing the damage caused by plant pathogens. Hence, plant breeders are continuously working to discover novel sources of genetic resistance. Crop wild relatives (CWRs) are a reservoir of genetic variation providing an important source of novel alleles for the genetic improvement of cultivated species. Crosses between cultivars and CWRs have been carried out in several crop species to unlock this favorable genetic diversity (Tanksley and McCouch, 1997; Feuillet et al., 2008). Some prominent examples of the introgression of favorable disease resistance alleles from CWRs are the introductions of late blight resistance into potato from the wild potato *Solanum demissum* (Rao, 1979; Prescott-Allen and Prescott-Allen, 1986), and of stem rust resistance genes *Sr21* (Chen et al., 2015) and *Sr39* (Kerber and Dyck, 1990), both effective against the race Ug99, into bread wheat from *Triticum monococcum* and *Aegilops speltoides*, respectively.

Barley (*Hordeum vulgare* L.), the fourth most important cereal crop in the world, is affected each year by up to 30% potential yield loss due to pests and diseases (Savary et al., 2012). Limitations in the availability of novel resistance genes or alleles in the primary gene pool of barley, comprising the cultivated barley *H. vulgare* spp. *vulgare* and its wild progenitor *H. vulgare* spp. *spontaneum*, has directed the focus of research towards other barley gene pools. Bulbous barley, *Hordeum bulbosum* (*Hb*), a wild self-incompatible species and the only member of the secondary gene pool of cultivated barley (von Bothmer et al., 1995) is resistant to many barley pathogens (Xu and Kasha, 1992; Walther et al., 2000). A large panel of *Hb* introgression lines (ILs) harboring a diverse spectrum of resistance traits has been developed during recent years (Pickering et al., 1995; Johnston et al., 2009). This resource comprises ILs carrying, among others, the barley leaf rust resistance gene *Rph26* on chromosome 1H^b^L (Yu et al., 2018); barley leaf rust gene *Rph22* (Johnston et al., 2013) and barley mild mosaic virus gene *Rym16*^*Hb*^ (Ruge-Wehling et al., 2006) both located on chromosome 2H^b^L; barley yellow dwarf virus resistance gene *Ryd4^Hb^* on chromosome 3H^b^L (Scholz et al., 2009) as well as loci conferring powdery mildew resistance located on chromosome 2H^b^S, 2H^b^L and 7H^b^L in barley/ *Hb* introgression lines (Xu and Kasha, 1992; Pickering et al., 1995; Shtaya et al., 2007). These ILs represent a unique genetic resource for improving barley resistance to pathogens and for scientific investigation of resistance mechanisms as they provide access to further genetic diversity out of the primary gene pool of barley (Tanksley and Nelson, 1996; Zamir, 2001; Johnston et al., 2009). Genotyping-by-sequencing (GBS) of 145 *Hv/Hb* introgression lines (Wendler et al., 2014, 2015) has provided an extensive pool of molecular markers, sequence resources and single-nucleotide polymorphisms (SNPs) information, greatly improving the efficiency of mapping *Hb* loci.

Since the identification and the extensive use of the durable and complete *mlo* resistance gene in European germplasm (Jørgensen, 1992), the threat of powdery mildew to barley has been largely mitigated. However, *mlo* is associated with yield penalties (Kjær et al., 1990) and increased susceptibility to some hemibiothrophic and necrotrophic fungi (Jarosch et al., 1999; Kumar et al., 2001; McGrann et al., 2014). Thus, the search for alternative sources of resistance to powdery mildew remains important for barley breeding (Czembor, 2002; Corrion and Day, 2015). Several *Hv*/*Hb* introgression lines carrying a locus conferring powdery mildew resistance have been described (Xu and Kasha, 1992; Pickering et al., 1995, 1998; Shtaya et al., 2007). The *Hb* accession A42 displays a dominant high resistance to powdery mildew that has been localized on the short arm of chromosome 2H^b^ in preliminary studies (Szigat and Szigat, 1991; Michel, 1996). Interestingly, several significant QTLs and major genes associated with powdery mildew resistance have repeatedly been reported in this region in cultivated barley (Backes et al., 2003; von Korff et al., 2005; Řepková et al., 2009). Moreover, a resistance gene to powdery mildew has been reported in this region in various *Hb* accessions (Pickering et al., 1995; Zhang et al., 2001; Shtaya et al., 2007).

While the potential value of untapped genetic diversity of CWR is immense, their application in breeding programs through the use of ILs is hampered by negative linkage drag, mainly caused by severely repressed genetic recombination (Wijnker and de Jong, 2008; Prohens et al., 2017), which can confer yield penalties or other unfavorable characteristics (Hospital, 2001). The degree of drag is correlated with the size of introgressed CWR chromatin segments, and thus can be mitigated by reducing the size of respective introgressions through recombination (Frisch and Melchinger, 2001). However, the efficient utilization of *Hb* germplasm in barley crop improvement and the genetic mapping of loci contributed by *Hb* suffers from highly reduced frequency of recombination in introgressed intervals up to 14-fold compared to intraspecific barley crosses (Ruge et al., 2003; Ruge-Wehling et al., 2006; Kakeda et al., 2008; Johnston et al., 2013). Possible explanations for this phenomenon include excessive sequence diversity, structural variation among *Hordeum* genomes, and probably other unknown mechanisms (Pickering, 1991; Hoffmann et al., 2004; Wendler et al., 2017). To reduce the negative linkage drag, precise delimitation of the causal gene is required, which usually demands intensive screening of large segregating *Hv/Hb* ILs populations.

Canady et al. (2006) compared the recombination rate in crosses between cultivated tomato (*Solanum lycopersicum*) and ILs with *Solanum lycopersicoides* fragments in *S. lycopersicum* backgrounds with the recombination in crosses between ILs with *S. lycopersicoides* fragments and ILs with *Solanum pennellii* fragments in the same region. They showed that tomato ILs with overlapping fragments from closely related species exhibited increased recombination rates in those fragments. Similarly, in barley, Johnston et al. (2015) demonstrated the usefulness of intraspecific recombination between a *Hb* ILs to overcome the negative linkage between genes conferring pathogen resistance and reduced yield. They crossed two recombinant ILs containing an *Hb* locus on chromosome 2HL comprising the genes *Rph22* and *Rym16^Hb^*, together with the proximal region of the original introgression for one of them, and the distal region for the other. They obtained four lines with reduced introgressions around the locus of interest for which the yield was close to the one of the recurrent *Hv* genotype.

The current study aimed to map a dominant powdery mildew resistance locus on chromosome 2H^b^S introgressed from the tetraploid A42 powdery mildew resistant *Hb* accession into a susceptible barley cultivar “Borwina”. Mapping in populations of over 200 F7 and BC1F6 from crosses between a susceptible barley cultivar and an *Hb* IL showed severely repressed recombination between the introgressed segment and the *Hv* genome. To overcome this difficulty, we exploited intraspecific recombination instead of interspecific recombination by crossing the IL carrying the resistance locus with another *Hv/Hb* IL carrying a homologous *Hb* introgression without resistance loci.

## 2 Material and Methods

### 2.1 Fungal isolate and phenotyping test

The swiss powdery mildew field isolate CH4.8 was chosen based on its very low infection rate on resistant plants (less than 5% of leaf area covered by colonies), allowing a clear discrimination between resistant and susceptible plants. The resistance test was carried out in two technical replicates by detached leaf assay as described in Hoseinzadeh et al. (2019). The inoculated detached leaves were kept for 7 days in a growth cabinet (MLR-352H, Panasonic, Japan) with LED light sources (S2 20W matt nw, ARTEKO-LED, Germany) at 20°C with 60% humidity and a 16:8 hours photoperiod. Powdery mildew resistance phenotype was scored macroscopically based on percentage infection area as described by Kølster et al. (1986) and Mains and Diktz (1930). Plants with low colony numbers and sizes surrounded by chlorotic and/or necrotic tissue (less than 25% leaf infection area) were classified as resistant, while those having infection types similar to the susceptible parent “Borwina”, with a large infection area covered by medium to large colonies with or without surrounding chlorosis, were classified as susceptible.

### 2.2 Plant material and population development

Initial genetic mapping was performed on the F7 and BC1F6 populations “5216” and “4176”, respectively, both derived from crossing a tetraploid derivative of the colchicine treated barley cultivar “Borwina” and the tetraploid *Hb* accession A42, which is resistant to several European barley powdery mildew isolates. Population “5216” is a recombinant inbred line (RIL) population whereas “4176” is a backcrossed inbred line (BCIL) population. The crossing scheme used to generate those populations is described in Figure 1.

**Figure 1:**
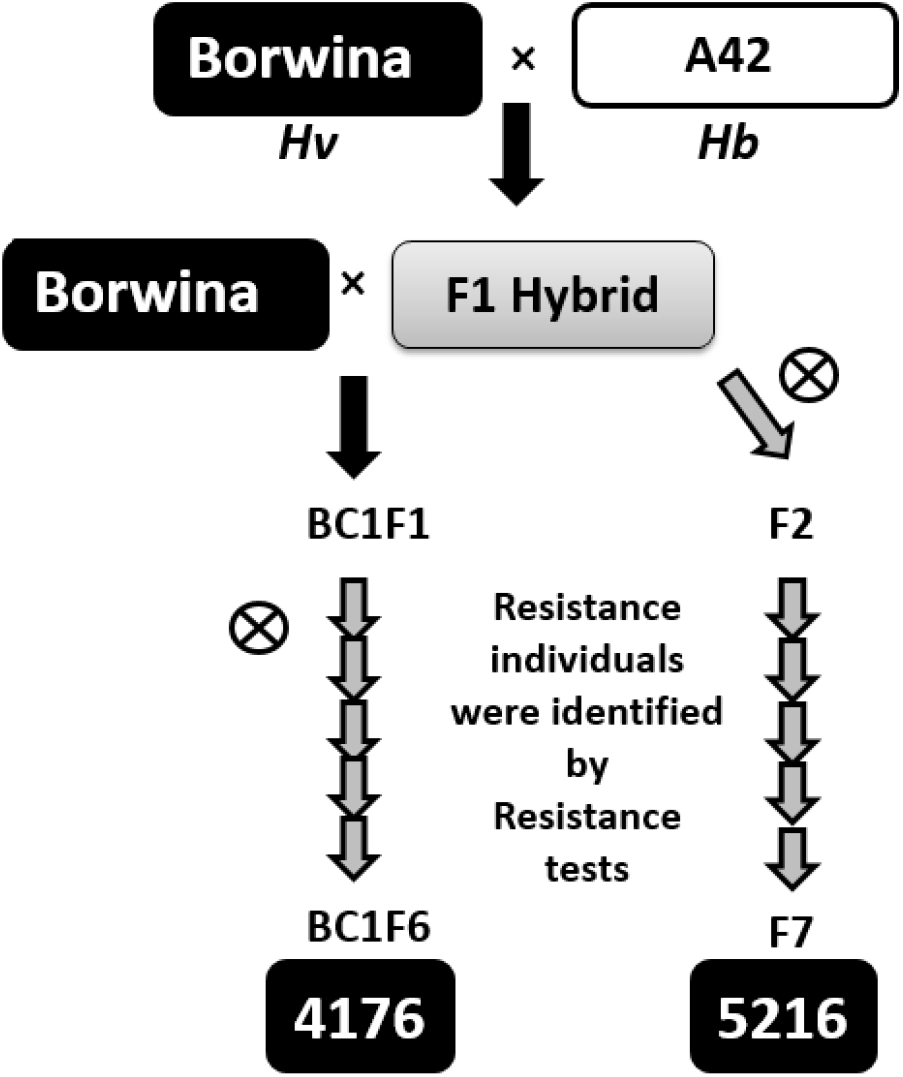
Schematic outline of the mapping population design. The tetraploid *Hb* accession A42, resistant to the isolate CH4.8, was crossed to the tetraploid derivative of the susceptible barley cultivar “Borwina”. From tetraploid F1 hybrids, two different introgression mapping populations were developed: the BCIL “4176” was developed by backcrossing the F1 hybrid once to the parent “Borwina”, followed by five generations of selfing. A grey arrow indicates a selfing generation. The RIL population, “5216” was derived from six generations of selfing from the F1 hybrid. In the course of development of populations “4176” and “5216”, the generations were diploid from BC1F2 and F2, respectively. Through the population development, the resistance test with the powdery mildew isolate CH4.8 and marker data analysis were performed to identify the resistant heterozygous IL to be selfed.

We hypothesized that differences in the sequence or organization between *Hv* and *Hb* orthologous genome regions would severely reduce meiotic recombination. To test this hypothesis, we analyzed intraspecific *Hb*/*Hb* recombination in a *Hv* background. We generated two F2 populations by crossing two independent *Hv/Hb* ILs carrying independent but overlapping *Hb* introgressions at the terminal 2HS chromosome, thus representing different *Hb* genotypes at the resistance locus. Three introgression lines “IL 88”, “IL 99” and “IL 116”, developed in New Zealand Institute for Crop and Food Research, were selected that carry independent *Hb* introgressions at end of the short arm of barley chromosome 2H (Wendler et al., 2015, Table 1). These three ILs were phenotyped for resistance against isolate CH4.8 (Figure 2). Only “IL 88” displayed full susceptibility to CH4.8 and was used for further population development.

**Table 1:** *Hb* introgression lines containing segments overlapping with IL “4176” and IL “5216” and their observed resistance phenotype to the *Bgh* isolate CH4.8.

| IPK ID | IL code (New Zealand) | Crossing scheme | introgression location GBS | Start (Mb) | End (Mb) | Phenotype of ILs to isolate CH4.8 |
| --- | --- | --- | --- | --- | --- | --- |
| 88 | 200A3/7/M1 | Emir x A17/1 | 2HS | 0 | 19,4 | Susceptible |
| 99 | 213G3/2/2/2/1 | Emir x (2920/4 x Tinos16/1) | 2HS, 6HS | 0 | 22,1 | Resistant |
| 116 | 230H24/5/M1/M1 | Morex x 2032 | 2HS | 0 | 19,4 | Resistant |

**Figure 2:**
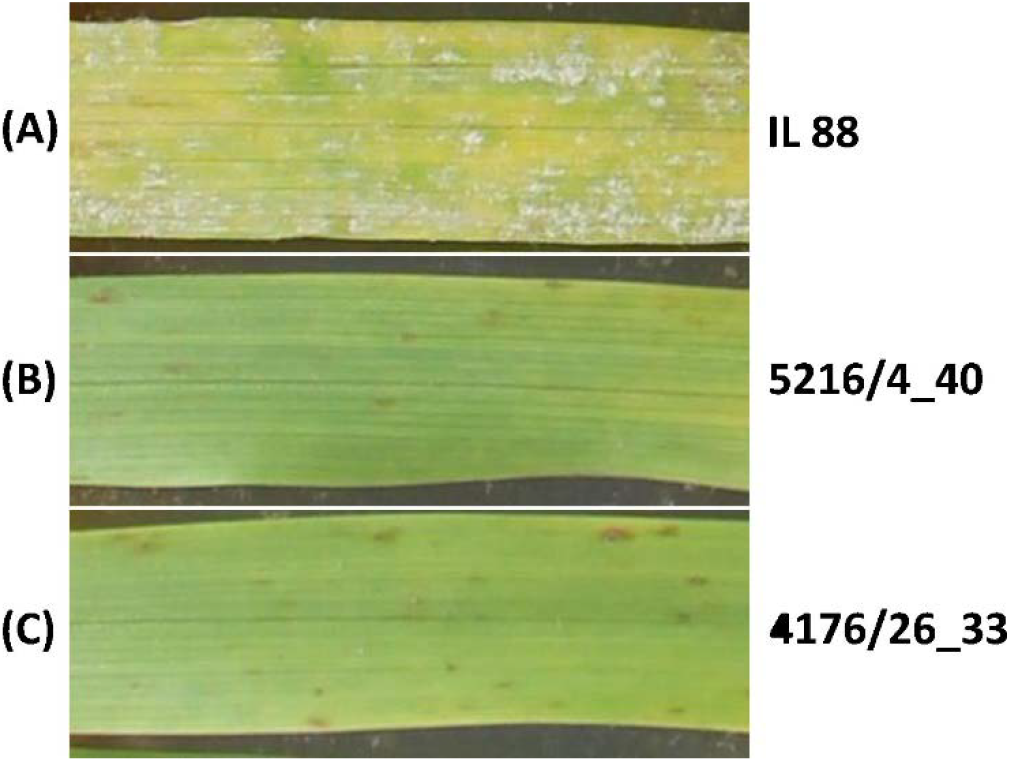
Powdery mildew infection (isolate CH4.8) on the second leaf of seedling from the three independent *Hv*/*Hb* parental ILs, seven days after inoculation. **(A)** Phenotype of the susceptible parental introgression line “IL 88” population “dIL_5216” and “dIL_4176”. **(B)** Phenotype of the resistant parental introgression line 5216/4_40 from population “dIL_5216”. **(C)** Phenotype of the resistant parental introgression line 4176/26_33 from population “dIL_4176”

The homozygous resistant ILs 5216/4_40 and 4176/26_33 from the populations “5216” and “4176” were crossed to the susceptible “IL 88”. From each cross, a single F1 plant was selfed, resulting in 103 and 146 F2 seeds, respectively. These two F2 populations were named “dIL_5216” and “dIL_4176”, respectively.

### 2.3 Genomic DNA extraction

Genomic DNA was extracted from third leaves of barley seedlings using a guanidine isothiocyanate-based DNA isolation method in 96-well plate format as described earlier (Milner et al., 2018). dsDNA concentration was measured by Qubit® 2.0 Fluorometer using the Qubit™ dsDNA BR (Broad Range) Assay Kit (Invitrogen, Carlsbad, CA, United States) following the manufacturer’s protocol.

### 2.4 Marker development

To screen the initial IL mapping populations “4176” and “5216” for recombinants, nine CAPS markers (supplementary Table 1) were designed to be evenly distributed over the distal 20 cM of barley chromosome 2HS, based on the conserved interspecific SNPs identified by targeted enrichment re-sequencing of 145 *Hb* ILs (Wendler et al., 2015). This set of ILs included the F4 homozygous 2HS IL (3026) from the mapping population “5216” and its associated donor lines, plus four additional *Hv* cultivars and four *Hb* accessions (Wendler et al., 2015). Only conserved *Hv/Hb* SNPs with a minimum of six-fold read coverage and located in the target region on the barley draft genome (International Barley Genome Sequencing Consortium, 2012) were selected and converted into CAPS markers using SNP2CAPS software (Thiel et al., 2004). Primer design was carried out using the default settings of Primer3 v.0.4.01 (Koressaar and Remm, 2007; O’Halloran, 2015) with minor modifications: The primer length was set between 19-21 bp. Primer melting temperature (Tm) was set to minimum Tm = 58°C, optimum Tm = 59°C and maximum Tm = 60°C. The product size was defined to be between 700 and 1,000 bp and Guanine-Cytosine content (GC-content) was set within the range of 50–55%.

### 2.5 CAPS genotyping

Genotyping of populations “4176” and “5216” with the described CAPS markers was performed in a 20 µl PCR reaction volume including 20 ng genomic DNA, 0.1 U of HotStarTaq DNA Polymerase (Qiagen, Hilden, Germany), 1x PCR reaction buffer containing 15 mM MgCl_2_ (Qiagen, Hilden, Germany), 0.2 mM of each dNTP (Fermentas, Fermentas, St. Leon-Rot, Germany), and 0.5 mM of each primer. All fragments were amplified using the following touchdown PCR profile: an initial denaturing step of 15 min at 95°C was followed by four cycles with denaturation at 95°C for 30 s, annealing at 62°C for 30 s (decreasing by 1°C per cycle), and extension at 72°C for 30 s. A final extension step was performed at 72°C for 7 min. The enzymatic digestion of the amplicons was performed in a 10 μl volume containing 5 μl of PCR product, 1× of appropriate buffer (New England Biolabs, Hitchin, UK), 1 U of enzyme (New England Biolabs, Hitchin, UK) and adjusted to final volume by adding molecular biology grade pure water. The reaction mix was incubated for one hour at recommended incubation temperature. The digested PCR products were resolved by 1.5–2.5 % gel-electrophoresis depending on amplicon size.

### 2.6 Genotyping-by-sequencing and data analysis

GBS was used, following published procedures (Mascher et al., 2013b), to check the genetic purity and state of heterozygosity of F1 hybrid seeds derived from crosses between overlapping 2HS introgression lines as well as to genotype the whole F2 populations derived from these crosses. DNA of the progeny and parental lines were pooled per Illumina HiSeq2500 lane in an equimolar manner and sequenced for 107 cycles, single read, using a custom sequencing primer as previously described (Hoseinzadeh et al., 2019). The reads were aligned to the TRITEX genome assembly of barley cultivar Morex (Monat et al., 2019) with BWA-MEM version 0.7.12a (Li and Durbin, 2009). Variants were filtered following the protocol of Milner et al. (2018) for a minimum depth of sequencing of four to accept a genotype call, a minimum mapping quality score of the SNPs (based on read depth ratios calculated from the total read depth and depth of the alternative allele) of six, a maximum fraction of heterozygous call of 70% and a maximum fraction of 25% of missing data. The resulting tables of polymorphisms are provided in supplementary tables 2 and 3.

## 3 Results

### 3.1 Inheritance of the resistance contributed by *Hb*

The susceptible parental *Hv* genotype “Borwina” consistently displayed a leaf infection area of ≥ 80% in all experiments. The scoring rates of all susceptible individuals of both populations “4176” and “5216” was similar to the susceptible parent and did not significantly vary between phenotyping experiments, indicating high infection efficiency and reproducibility in all phenotyping experiments. The resistance phenotype to the CH4.8 powdery mildew isolate of plants carrying the *Hb* introgressed segment in a heterozygous state was identical to that of plants homozygous for the *Hb* fragment (Figure 3). The resistance phenotype was invariably accompanied with a hypersensitive response (HR) forming a necrotic lesion.

**Figure 3:**
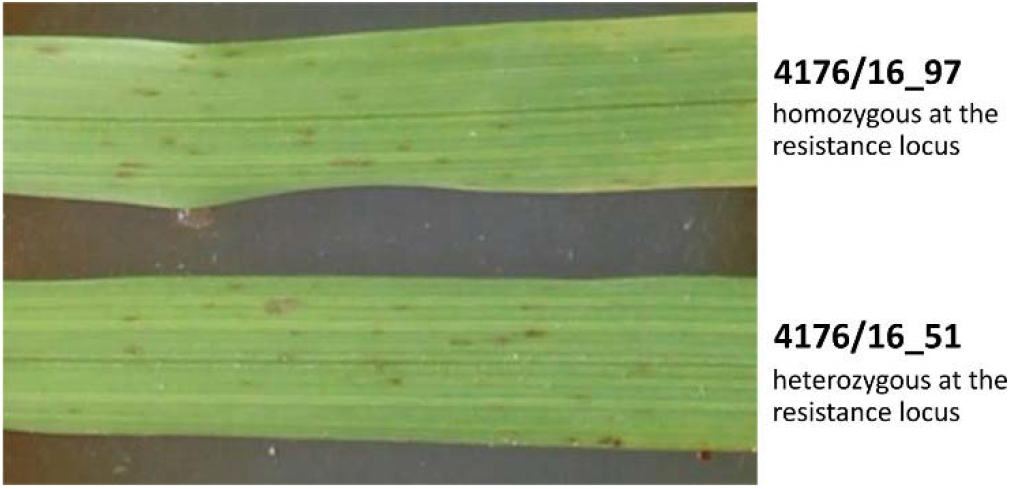
Powdery mildew infection (isolate CH4.8) on the second leaf of seedling from two BC2F6 ILs from population “4176”, seven days after inoculation. 4176/16_97 is homozygous at the resistance locus whereas 4176/16_51 is heterozygous. Their resistance phenotype is identical and present necrotic lesions characteristic of HR.

Phenotypic segregation for powdery mildew resistance against CH4.8 isolate was consistent with a 3:1 ratio (resistant/susceptible, R/S, P<0.05) in all mapping populations, indicating the control of resistance by a single dominant resistance gene (Table 2). We propose the temporary name *Mlhb.A42* for this locus, based on previous naming of *Hb* powdery mildew resistance genes (Pickering et al., 1995; Steffenson, 1998).

**Table 2:**
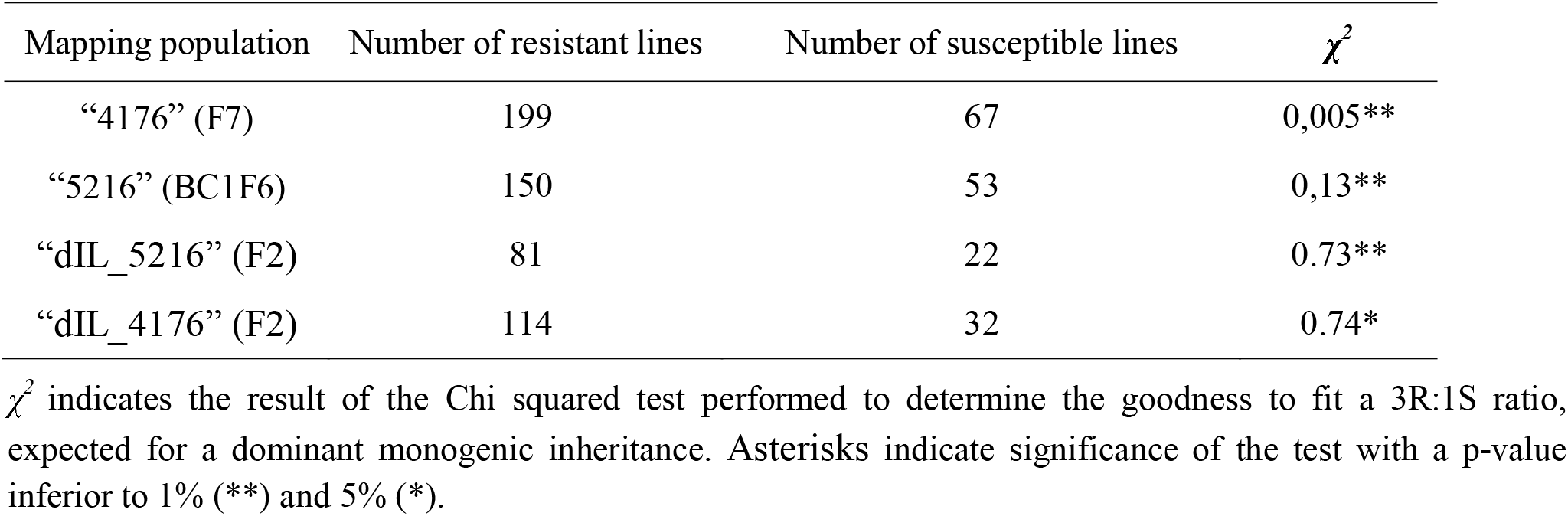
Phenotypic segregation of powdery mildew resistance in each of 2HS IL mapping populations.

### 3.2 Recombination frequency and mapping of *Mlhb.A42* in single-introgression line mapping populations

Genotyping of the mapping populations employed nine CAPS markers designed based on existing exome capture re-sequencing data (Supplementary table 1). This showed that the BC1F6 population “4176” carried a longer introgressed segment compared to the F7 population “5612” (Figure 4). The genotyping results for population “5612” confirmed the extent of the introgressed segment, previously detected by exome capture in the F6 generation (Wendler et al., 2015).

**Figure 4:**
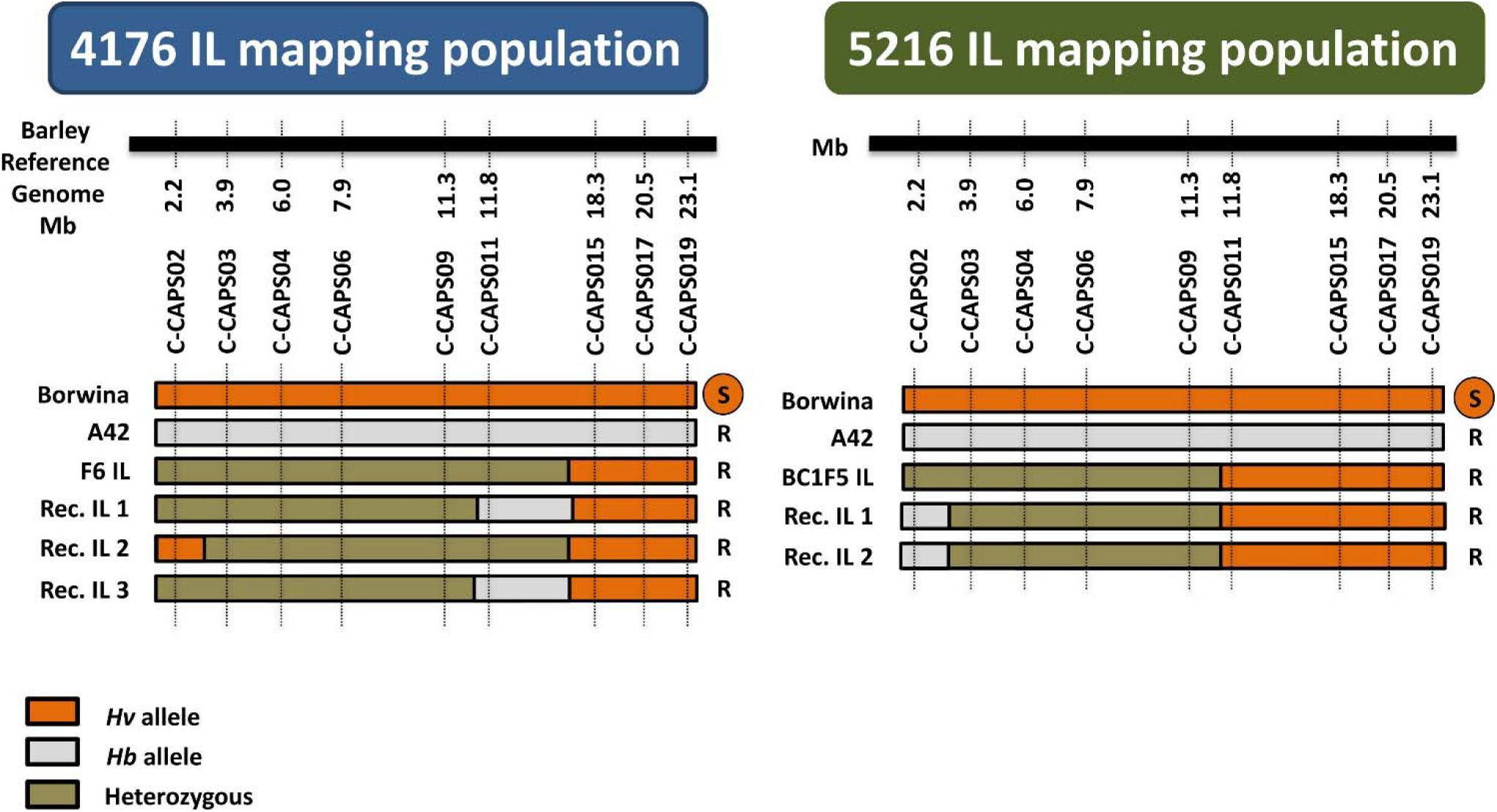
Graphical genotypes and phenotypes of the recombinant *Hv/Hb* ILs from the two IL mapping populations. **“**Borwina” and “A42” are the original susceptible and resistant parents of the populations. “F6 IL” and “BC1F5 IL” represent the plant that was selfed in the previous generation to obtain populations “4176” and “5216”, respectively. “Rec. IL” 1 to 3 are the identified recombinant plants. Nine CAPS markers named C-CAPS02 to C-CAPS019, were developed based on conserved *Hv/Hb* SNP loci described in Wendler et al (2014), spanning the terminal 23 Mbp of barley chromosome 2HS. The black horizontal bars represent schematically the barley reference genome. The physical position of each selected SNPs on the barley reference genome is written below the black line. The homozygous state of the alleles from susceptible and resistant parents is shown as orange and grey colors, respectively, whereas heterozygous state is shown in green. The phenotype of each recombinant is indicated on the right of their genotype (R = resistant; S = susceptible). The number of recombinants identified through screening with the developed markers was only three and two in IL mapping population “4176” and “5216”, respectively.

Genotyping of 266 and 203 individuals in the initial IL mapping populations “4176” and “5612”, uncovered only three and two recombinants within the introgressed segments, respectively (Figure 4). The introgressed segment therefore has a genetic length of approximately 1 centiMorgan (cM). Yet, the segment of population “5612” represents 10 cM on the barley POPSEQ map (Mascher et al., 2013a; Wendler et al., 2015). This confirms the anticipated reduced recombination between the *Hv* genome and the introgressed *Hb* segment. The resulting genetic interval for *Mlhb.A42* is flanked by markers CAPS02 and CAPS11, corresponding to a 9.5 Mbp interval between bp positions 2,269,761 and 11,819,688 on chromosome 2HS.

### 3.3 Recombination and mapping of *Mlhb.A42* in double introgression lines

The 103 and 146 F2 plants from the two double introgression populations “dIL_5216” and “dIL_4176”, respectively, were genotyped by GBS and phenotyped for resistance against CH4.8 isolate. Based on obtained SNP variants across candidate interval for *Mlhb.A42* defined in the initial mapping populations, 14 and 36 recombinants were obtained in “dIL_5216” and “dIL_4176”, respectively (Figure 5). The phenotypes of all non-recombinant plants corresponded to their genotype in this interval. Association between the phenotypes and genotypes of the recombinants defined overlapping 1.7 Mbp (between 7,943,409 and 9,595,691 bp) and 1.4 Mbp (between 8,193,151 and 9,595,691 bp) intervals on barley chromosome 2HS (Monat et al., 2019) in “dIL_5216” and “dIL_4176”, respectively (Figure 5).

**Figure 5:**
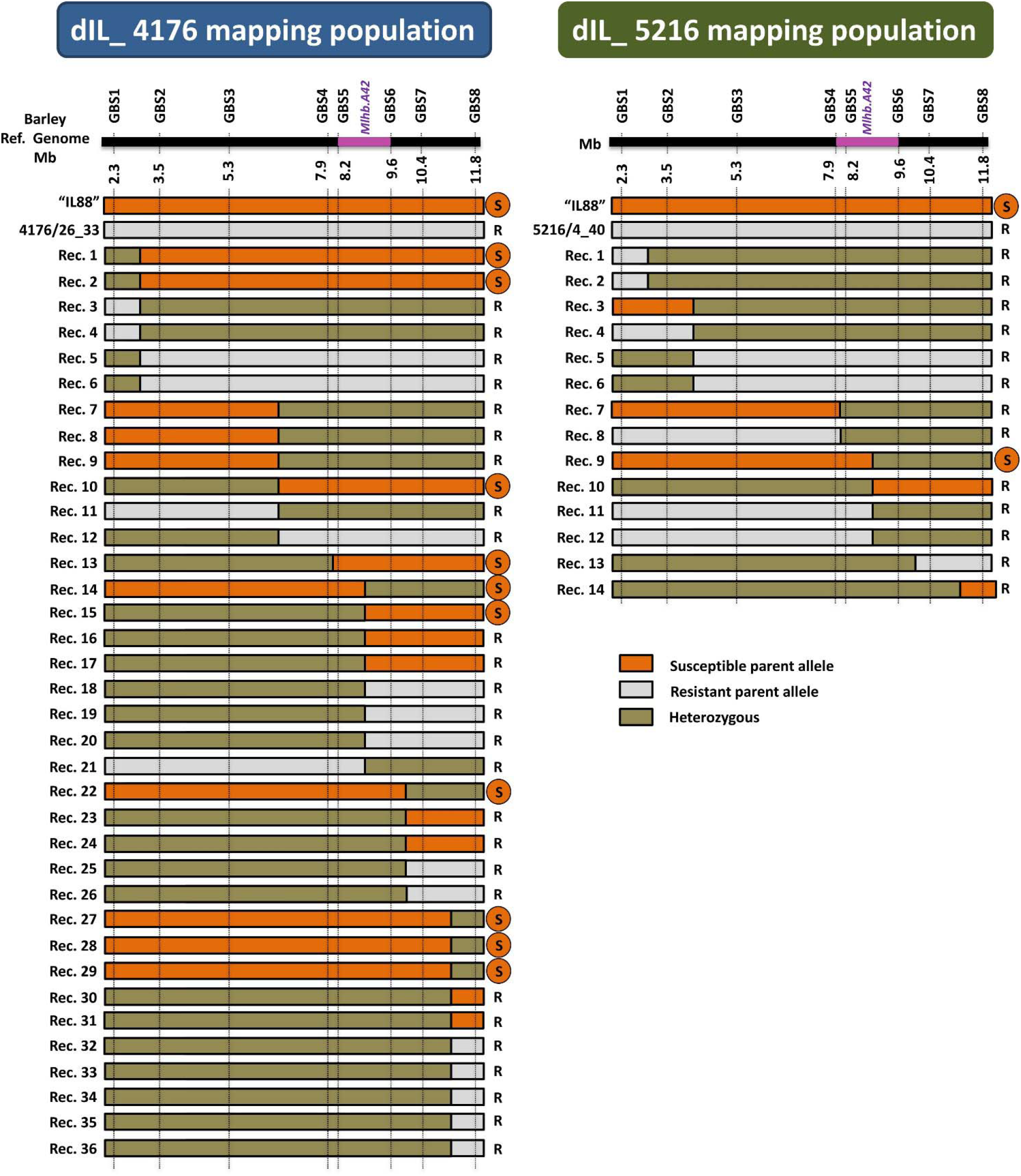
Intraspecific mapping of the powdery mildew resistance locus *Mlhb.A42* in F2 populations derived from two different 2HS *Hv*/*Hb* introgression lines. The horizontal black bar schematically represents barley chromosome 2HS. The physical coordinates of each GBS marker are indicated below it. The genomic region containing the *Mlhb.A42* locus on barley chromosome 2HS is shown in purple. The graphical genotypes of recombinants are indicated as horizontal bars. The genotype of the susceptible parent “IL88” is represented in orange, the one of the resistant parents 4176/26_33 and 5216/4_40 is represented in grey, and heterozygous state is shown in olive green. The phenotype of each recombinant plant is indicated on the right of its genotype (R= resistant; S = susceptible). The recombination rates in “dIL_4176” and “dIL_5216” are 24.7 % and 13.6%, respectively, corresponding to a 20 fold increase compared to single introgression lines. The *Mlhb.A42* gene was allocated to a 1.4 Mb interval, based on an estimate using the *Hv* genome as reference.

### 3.4 Candidate genes within the target interval

Based on the annotated barley reference genome sequence (Monat et al., 2019), 26 high confidence (HC) genes (Table 3) are located within the 1.4 Mbp *Mlhb.A42* interval as delimited in the “dIL_4176” population. Those genes with a putative functional annotation included an HR-like lesion-inducing protein, a lectin receptor kinase (LecRK) and a Heat shock protein 90, all gene functions that could be directly or indirectly implied with plant resistance to biotic and/or abiotic stresses and therefore, qualify as candidate genes for *Mlhb.A42*. HORVU.MOREX.r2.2HG0082250 is annotated as a LecRK. However, analysis of its protein sequence with InterPro (Mitchell et al., 2018), showed that, like LecRKs, it is composed of an extracellular legume (L-type) lectin domain and a transmembrane domain. However, its cytoplasmic domain is only 23 amino-acid long and does not bear a kinase domain, making this gene a probable lectin receptor-like protein (LecRLP).

**Table 3:** High confidence (HC) genes based on automated annotation of barley reference genome

| Gene name | Start<sup>1</sup> | End<sup>1</sup> | Annotation |
| --- | --- | --- | --- |
| HORVU.MOREX.r2.2HG0081680 | 8302525 | 8304486 | 2-oxoglutarate (2OG) and Fe(II)-dependent oxygenase superfamily protein, putative |
| HORVU.MOREX.r2.2HG0081720 | 8364720 | 8365206 | NADH-ubiquinone oxidoreductase chain 1 |
| HORVU.MOREX.r2.2HG0081740 | 8381983 | 8383512 | ATP synthase subunit alpha |
| HORVU.MOREX.r2.2HG0081760 | 8470715 | 8473118 | HR-like lesion-inducing protein-related protein |
| HORVU.MOREX.r2.2HG0081770 | 8474573 | 8487616 | Actin-related protein |
| HORVU.MOREX.r2.2HG0081780 | 8511271 | 8512941 | Cytochrome P450 |
| HORVU.MOREX.r2.2HG0081810 | 8576680 | 8577147 | Cytochrome P450 |
| HORVU.MOREX.r2.2HG0081820 | 8607940 | 8608750 | Cytochrome P450 |
| HORVU.MOREX.r2.2HG0081830 | 8615323 | 8617046 | Cytochrome P450 |
| HORVU.MOREX.r2.2HG0081840 | 8664852 | 8679924 | Cytochrome P450 |
| HORVU.MOREX.r2.2HG0081850 | 8686994 | 8691803 | Kaurene synthase |
| HORVU.MOREX.r2.2HG0081880 | 8896440 | 8898414 | Copalyl diphosphate synthase |
| HORVU.MOREX.r2.2HG0081920 | 8961925 | 8964012 | Copalyl diphosphate synthase |
| HORVU.MOREX.r2.2HG0082000 | 9064624 | 9068933 | Copalyl diphosphate synthase |
| HORVU.MOREX.r2.2HG0082020 | 9253167 | 9258032 | Agenet domain, putative |
| HORVU.MOREX.r2.2HG0082060 | 9337837 | 9339296 | O-methyltransferase family protein |
| HORVU.MOREX.r2.2HG0082080 | 9357711 | 9359256 | Glycosyltransferase |
| HORVU.MOREX.r2.2HG0082140 | 9435844 | 9439807 | Transcription factor |
| HORVU.MOREX.r2.2HG0082180 | 9468781 | 9469636 | F-box protein PP2-A13 |
| HORVU.MOREX.r2.2HG0082190 | 9472177 | 9476645 | ATP sulfurylase (Sulfate adenylyltransferase) |
| HORVU.MOREX.r2.2HG0082200 | 9478069 | 9482961 | Zinc finger family protein |
| HORVU.MOREX.r2.2HG0082240 | 9505214 | 9505813 | Maternal effect embryo arrest protein |
| HORVU.MOREX.r2.2HG0082250 | 9538406 | 9539521 | Lectin receptor kinase |
| HORVU.MOREX.r2.2HG0082260 | 9543886 | 9546930 | Carboxymethylenebutenolidase-like protein |
| HORVU.MOREX.r2.2HG0082310 | 9591121 | 9593401 | Heat shock protein 90 |
| HORVU.MOREX.r2.2HG0082320 | 9595584 | 9597283 | caspase-6 protein |

<sup>1</sup>Coordinates based on TRITEX Morex assembly (Monat et al., 2019).

## 4 Discussion

We report the fine mapping of *Mlhb.A42*, a dominant powdery mildew resistance locus originating from *Hb*, using mapping populations derived by crossing two independent, resistant and susceptible, *Hv*/*Hb* ILs. The strong suppression of recombination between homeologous genomic segments in *Hv*/*Hb* introgression lines, which typically results in severe linkage drag, previously represented a barrier to the efficient utilization of *Hb* germplasm in barley crop improvement, and to the isolation of disease resistance genes introgressed from *Hb*.

The genomic resource created by the GBS study of Wendler et al. (2015) on 145 *Hv/Hb* ILs proved to be a useful tool to identify suitable partners for the development of double ILs populations. The exploitation of intraspecific recombination allowed us to overcome the barrier to recombination usually observed in IL populations. Recombination rates in the region carrying the introgressed fragment were estimated to be 24.7 % (“dIL_4176”) and 13.6 % (“dIL_5216”), comparable to the corresponding 10 % rate observed in pure *Hv/Hv* mapping populations (e.g. POPSEQ map (Mascher et al., 2013a)). The rate of recombination was exceeded by a factor of 20-fold as compared to the F7 and BC1F6 single IL populations “5216” and “4176” (approximately 1%). The polymorphisms and markers identified in this study can be converted into KASP assays which would enable for rapid and high-throughput screening of large breeding population for the purpose of introgression of *Mlhb.A42* into barley cultivars.

The 1.4 Mbp identified target interval is containing 26 annotated HC genes on the most recent chromosome-scale genome assembly of cultivar ‘Morex’ (Monat et al., 2019). The powdery mildew resistance conferred by this *Hb* locus is dominantly inherited, displaying chlorotic/necrotic flecks of HR with collapsed hyphae at inoculation sites, suggesting a salicylic acid (SA)-independent resistance pathway. Genes from the nucleotide-binding-site leucine-rich-repeat (NBS-LLR) or the receptor-like-kinase (RLK) families are over-represented among genes conferring this type of strong dominant resistance to pathogens (Kourelis and van der Hoorn, 2018), making genes from these families likely candidates in the context of this study. None of the annotated genes in the interval are part of the NBS-LRR family; however HORVU.MOREX.r2.2HG0082250 is annotated as a LecRK. LecRKs are a type of RLK characterized by an extracellular lectin motif (Wu and Zhou, 2013). They have been described as implicated in biotic stress resistance, mostly to bacteria and fungi (Singh and Zimmerli, 2013). This type of gene has been identified in resistance to oomycetes (Wang et al., 2015b, 2015a; Balagué et al., 2017) and fungi (Huang et al., 2013; Wang et al., 2014a) in *Arabidopsis thaliana* and to wheat powdery mildew in *Haynaldia villosa* (Wang et al., 2018). According to InterPro (Mitchell et al., 2018), HORVU.MOREX.r2.2HG0082250 does not bear a kinase domain and is likely to be a L-type LecRLP. So far, the only LecRLPs described have a lectin-like Lysin-motif (LysM)-type lectin domain. Two LysM-LecRLPs from *A. thaliana* and three from rice have been identified reported in context of disease resistance through interaction with the LysM-LecRK CERK1. The rice LysM-LecRLP CEBiP recognizes chitin and, through a direct interaction with CERK1, confers pattern-triggered immunity against fungi (Shimizu et al., 2010). Similarly, LYP4 and LYP6 both perceive peptidoglycan and chitin and interact with CERK1 (Liu et al., 2012). In *A. thaliana*, LYM1 and LYM3 sense pectidoglycan and trigger immunity, through CERK1, to bacterial infection (Willmann et al., 2011). *A. thaliana* contains only four L-type LecRLPs (Bellande et al., 2017) but, so far, no L-type LecRLPs have been functionally described. However, a mode of action of HORVU.MOREX.r2.2HG0082250 similar to the one of LysM-LecRLPs is a possibility for further investigation.

Almost all the genes annotated in the delimited target interval might be directly or indirectly involved in resistance to biotic and abiotic stresses and should be considered tentative candidates. NADH-ubiquinone oxidoreductase is involved in intracellular ROS production that can prevent pathogen infection (van der Merwe and Dubery, 2006). Cytochrome P450 and O-methyltransferase proteins are responsible for production of several molecules that can play a role in resistance to pathogens or pests (He and Dixon, 2000; Dixon, 2001; Noordermeer et al., 2001). Some glycosyl-transferase genes have been identified as necessary for the HR (Langlois-Meurinne et al., 2005). The rice gene *OsHRL* encodes for a HR-like lesion inducing protein and has been shown to be associated with resistance to bacterial blight (Park et al., 2010). Copalyl-diphosphate synthases are implicated in the biosynthesis of phytohormones including gibberellic acid or phytoalexins, which contribute to defensive secondary metabolism (Prisic et al., 2004; Harris et al., 2005). Finally, HORVU.MOREX.r2.2HG0082310 is annotated as a Heat shock protein 90 (Hsp90), which are involved in stress resistance, in particular disease resistance, mediating signal transduction for HR (Xu et al., 2012). Notably, it has been shown by virus-induced gene silencing that an *Hsp90* gene is required for *Mla13* resistance of barley to powdery mildew (Hein et al., 2005).

In addition to the annotated genes, other genes, e.g. members of NBS-LRR and RLK families, might be present in the interval of the resistant *Hb* parent but missing in the respective interval of the ‘Morex’ reference sequence. Indeed, *Hv* and *Hb* diverged 6 million years ago, accumulating structural variations since then (Jakob and Blattner, 2006). Therefore, this *Hb* resistance to powdery mildew could be due to a gene absent from the barley reference genome. In particular, NBS-LRR genes are subject to frequent duplication (Flagel and Wendel, 2009), and resistances conferred by NBS-LRRs genes are frequently due to presence/absence variation of such genes (Grant et al., 1998; Griffiths et al., 1999; Henk et al., 1999; Stahl et al., 1999; Tian et al., 2002). The wheat powdery mildew-resistance gene *Pm21* (Xing et al., 2018) originates from the wheat/*Dasypyrum villosum* translocation line T6AL.6VS and is localized in a region presenting a high level of synteny with the *Mlhb.A42* locus. This gene confers broad spectrum dominant resistance against wheat powdery mildew isolates and encodes a RPP13-like NBS-LRR gene (He et al., 2017; Xing et al., 2018). Its localization in a syntenic region to the resistance conferring *Mlhb.A42* locus provides a tempting working hypothesis that *Pm21* and *Mlhb.A42* could represent orthologous genes or members of the same locally evolved gene family.

A resequencing strategy of the *Mlhb.A42* locus in both resistant and susceptible *Hb* will be necessary to ascertain the structure of the locus and the number and nature of the candidate genes. With the development of new sequencing methods this could be achieved by Cas9-guided enrichment of the locus (Wang et al., 2014b) or sorting and sequencing for assembly of the chromosome 2H in one of the two introgression lines (Thind et al., 2017). Moreover, a locus of resistance to powdery mildew has been identified on chromosome 2HS in the *Hb* accessions S1 (Pickering et al., 1995; Shtaya et al., 2007), 2032 (Zhang et al., 2001; Shtaya et al., 2007), and A17 (Shtaya et al., 2007). Allelism tests are required to check whether the same locus is involved in the resistance from those four accessions or not.

Durability and spectrum of a resistance gene are two major criteria to assess its application potential (Mundt, 2014). Durability of resistance genes can become a major concern as deployed resistance genes are experiencing a boom-and-bust phenomenon (Gladieux et al., 2015). The spectrum of *Mlhb.A42* was not evaluated in this study and should be tested in order to ascertain its potential for field resistance. Durability is difficult to estimate under laboratory scenarios. The durability of orthologues genes can be used as a proxy, yet quite imperfect. As discussed earlier, the wheat resistance gene *Pm21*could be orthologous to *Mlhb.A42*. Varieties carrying *Pm21* have increasingly been cultivated in China in the recent years (Bie et al., 2015) and its durability can therefore be evaluated in real conditions. Unfortunately, in some wheat fields, new *Bgt* isolates, virulent against *Pm21* have been identified (Shi et al., 2009; Yang et al., 2009). However, resistance based on this gene persisted close to 40 years (Tang et al., 2018) and is still effective against more than a thousand of field isolates in China (Zeng et al., 2014) and Poland (Czembor et al., 2014). To counteract the risk of isolates breaking the resistance provided by a single locus it is of ongoing importance to identify new resistance genes and to pyramid new loci with existing sources of resistance to increase the durability of resistance in the system (Wu et al., 2019). Moreover, exploiting CWR resistances could be a way to unravel even more effective resistance genes. Indeed, Wang et al. (2019) showed that the *Hb* LecRLK gene of resistance to leaf rust *Rph22* confers a stronger resistance to leaf rust adapted to *Hv* than its *Hv* ortholog *Rphq2*. The hypothesis is that crop receptors have a lower recognition of crop-adapted pathogens than the CWR receptors because of adaptation of the pathogens during centuries of coevolution with their host plant.

In the current study we report genetic mapping of *Mlhb.A42*, a dominant resistance locus introgressed to cultivated barley from *Hb*. This work is a proof-of-concept study for establishing the basic steps of map-based cloning of genes present in *Hv/Hb* IL collections by exploiting double ILs mapping populations. Using this strategy, we circumvented the limitation of repressed meiotic recombination which was frequently observed in attempts of genetic mapping employing populations derived between *Hv*/*Hb* introgression lines and pure barley cultivars. Here, we observed similar or even higher recombination rates as expected in *Hv* and thus providing a major step towards facilitated exploitation of secondary gene pool-derived resistance genes in barley crop improvement.

## 6 Author Contributions

PH performed the experimental work. PH and HP performed data analysis and wrote the manuscript. BRW provided seed material and sample information for two IL mapping populations “4176” and “5216”. PS supervised the phenotyping. NS designed the study, supervised the experimental work, and contributed to the writing of the manuscript. All authors read, corrected, and approved the manuscript.

## 8 Acknowledgments

We gratefully appreciate the excellent technical support by Mary Ziems in performing crossings and Susanne Koenig in GBS library preparation. We kindly acknowledge Dr. Axel Himmelbach for his valuable support in the sequencing, Dr. Martin Mascher for his invaluable guidance in data processing and analysis, and Dr. Timothy Rabanus-Wallace for language editing. We are grateful to Dr. Neele Wendler for her support in the employment of the previously developed GBS and exome capture data presented in the current study.

## 12 Supplementary Material

Supplementary table 1: Characteristics of the CAPS markers developped for genotyping of populations "4176" and "5216".

Supplementary table 2: Variants in population "dIL_4176" as obtained after filtration of GBS data.

Supplementary table 3: Variants in population "dIL_5216" as obtained after filtration of GBS data.

